# Agmatine accumulation by *Pseudomonas aeruginosa* clinical isolates confers antibiotic tolerance and dampens host inflammation

**DOI:** 10.1101/299099

**Authors:** Jennifer L. McCurtain, Adam J. Gilbertsen, Clayton Evert, Bryan J. Williams, Ryan C. Hunter

## Abstract

In the cystic fibrosis (CF) airways, *Pseudomonas aeruginosa* undergoes diverse physiological changes in response to inflammation, antibiotic pressure, oxidative stress and a dynamic bioavailable nutrient pool. These include loss-of-function mutations that result in reduced virulence, altered metabolism and other phenotypes that are thought to confer a selective advantage for long-term persistence. Recently, clinical isolates of *P. aeruginosa* that hyperproduce agmatine (decarboxylated arginine) were cultured from individuals with CF. Sputum concentrations of this metabolite were also shown to correlate with disease severity. This raised the question of whether agmatine accumulation might also confer a selective advantage for *P. aeruginosa in vivo* during chronic colonization of the lung. Here, we screened a library of *P. aeruginosa* CF clinical isolates and found that ~5% of subjects harbored isolates with an agmatine hyperproducing phenotype. Agmatine accumulation was a direct result of mutations in *aguA*, encoding the arginine deiminase that catalyzes the conversion of agmatine into various polyamines. We also found that agmatine hyperproducing isolates (*aguA-*) had increased tolerance to the cationic antibiotics gentamicin, tobramycin and colistin relative to their chromosomally complemented strains (*aguA*+). Finally, we revealed that agmatine diminishes IL-8 production by airway epithelial cells in response to bacterial infection, with a consequent decrease in neutrophil recruitment to the murine airways in an acute pneumonia model. These data highlight a potential new role for bacterial-derived agmatine that may have important consequences for the long-term persistence of *P. aeruginosa* in the CF airways.

## INTRODUCTION

Cystic fibrosis (CF) is a lethal autosomal recessive disorder characterized by abnormal transepithelial ion transport and dehydrated mucus lining the epithelium of several organs, including the lung (1). Within the airways, compromised innate immunity and impaired mucociliary transport facilitate chronic colonization by variety of microorganisms that are the major cause of patient morbidity (2,3). Though recent culture-independent surveys have revealed complex CF lung microbiota consisting of hundreds of bacterial species (4–6), *P. aeruginosa* remains widely recognized as the primary driver of disease progression (7). This bacterium is prevalent among 70-80% of CF adults (8), can reach densities as high as 10^9^ cells/gm of sputum despite aggressive antimicrobial therapies (9), and its persistence strongly correlates with poor disease outcomes (10,11).

As chronic infections progress, *P. aeruginosa* is exposed to selective pressures that include the host immune response, competing microbiota, xenobiotics, oxidative stress and a dynamic nutritional milieu. In response, *P. aeruginosa* undergoes substantial genotypic and phenotypic variability. For example, loss-of-function mutations are commonly found in *lasR*, encoding a transcriptional regulator of quorum sensing (12). In turn, *P. aeruginosa* clinical isolates exhibit altered expression of QS-regulated virulence effector molecules such as elastase, siderophores and phenazines (13–16). Additional mutations in *vfr, exsA, mutS, ampR and mucA*, among others, lead to mucoidy, auxotrophy, altered motility, LPS modifications, hypermutability, and decreased susceptibility to phage, antimicrobials and phagocytosis (17–24). Each of these phenotypes is thought to confer a selective advantage for long-term persistence *in vivo* (25).

Recently, direct measurements of CF sputum have revealed elevated concentrations (>10µM) of agmatine, a pre-polyamine intermediate metabolite of the arginine decarboxylase pathway (Fig. 1)(26). *P. aeruginosa* isolates derived from CF subjects have also been found to accumulate this metabolite *in vitro*, though the genetic basis of this phenotype has not yet been determined (26). Previous studies have reported elevated concentrations of the polyamines spermidine and putrescine (that are derived from agmatine) in CF sputum and bronchoalveolar lavage fluid (27,28). Others have demonstrated that for *P. aeruginosa*, polyamines contribute to increased resistance to antimicrobials and oxidative stress (29). These observations raised the question: do loss-of-function mutations leading to agmatine hyperproduction also confer an advantage to *P. aeruginosa* in the context of CF lung infection?

**Figure 1.**
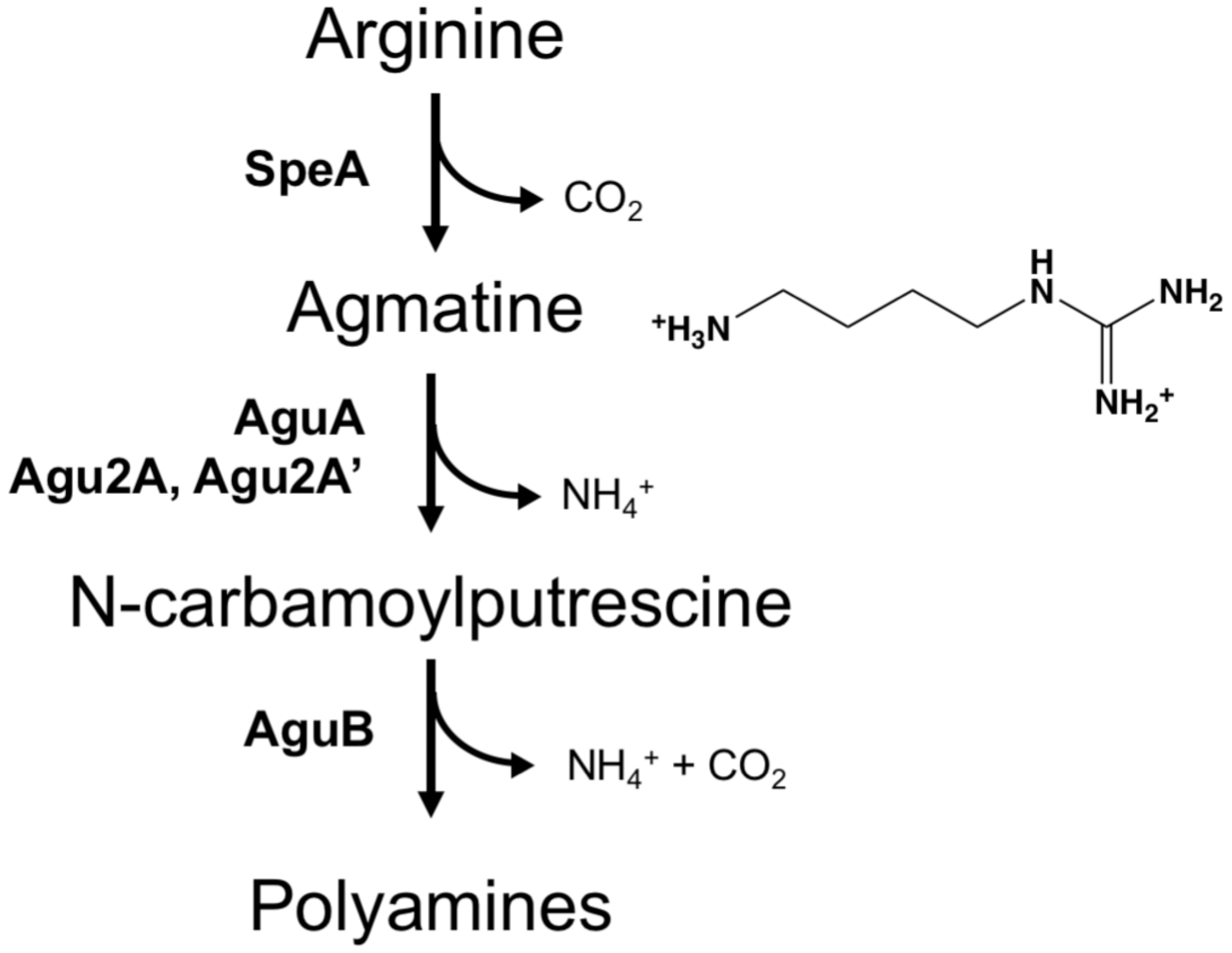
Arginine metabolism in *P. aeruginosa*. Agmatine is a pre-polyamine intermediate metabolite of the arginine decarboxylation pathway. Its hyperproduction results from mutations in *aguA*, encoding agmatine deiminase.

The objectives of this study were two-fold. First, we sought to characterize the frequency of agmatine hyperproducers among a library of *P. aeruginosa* CF isolates and to determine the genetic basis of this phenotype. Next, we tested whether agmatine accumulation protects *P. aeruginosa* against the host immune response and antibiotic stress. We demonstrate that agmatine confers a strain-dependent persistence advantage in the context of CF lung disease and highlight a potential new role for agmatine during *P. aeruginosa* infections of the lower airways.

## MATERIALS AND METHODS

### Bacterial strains, human epithelial cells and culture conditions

Bacterial strains and plasmids used in this study are presented in Table 1. Strains were routinely cultured at 37°C on Luria Bertani (LB) medium or Mueller Hinton Broth 2 (MHB-2) as indicated. When necessary, antibiotic concentrations were added as follows: for *Escherichia coli*, 20 µg/mL tetracycline, 50 µg/mL ampicillin; for *P. aeruginosa*, 25 µg/mL irgasan, 25-200 µg/mL tetracycline, and 200 µg/mL carbenicillin. Clinical isolates (one per subject) were derived from stable adult outpatients at Children’s Hospital Boston or the University of Minnesota. Studies were approved by the Committee on Clinical Investigation at CHB (#09-04-0183) and the Institutional Review Board at UMN (#1008E88194). Cystic fibrosis human bronchial epithelial cells (CF-DHBE) were acquired from Lonza and cultured using Bronchial Epithelial Cell Growth Medium (BEGM) and subculturing reagents (Lonza) according to the manufacturer’s protocol. Growth medium was changed every 48h until cells were ~80-90% confluent.

**Table 1.** Bacterial strains and plasmids.

| Parent Strain | Genotype / Phenotype | Source or Reference |
| --- | --- | --- |
| PA14 | Wildtype | 30 |
|  | Agmatine reporter, *aguA::gm Δagu2ABCA' ΔspeA aguRB-lux* | 31 |
|  | Agmatine deiminase knockout, *ΔaguAΔagu2ABCA'* | 32 |
| MNPA04 | Agmatine overproducing clinical isolate | This study |
|  | MNPA04 with intact arginine deiminase, *+aguA* | This study |
|  | MNPA04 with arginine decarboxylase knockout, *ΔspeA* | This study |
| MNPA05 | Agmatine overproducing clinical isolate | This study |
|  | MNPA05 with intact arginine deiminase, *+aguA* | This study |
|  | MNPA05 with arginine decarboxylase knockout, *ΔspeA* | This study |
| MNPA06 | Agmatine overproducing clinical isolate | This study |
|  | MNPA06 with intact arginine deiminase, *+aguA* | This study |
|  | MNPA06 with arginine decarboxylase knockout, *ΔspeA* | This study |
| CHB54-11 | Agmatine overproducing clinical isolate | This study |
|  | CHB54-11 with intact arginine deiminase, *+aguA* | This study |
|  | CHB54-11 with arginine decarboxylase knockout, *ΔspeA* | This study |
| CHB47-6 | Agmatine overproducing clinical isolate | This study |
|  | CHB47-6 with intact arginine deiminase, *+aguA* | This study |
|  | CHB47-6 with arginine decarboxylase knockout, *ΔspeA* | This study |
| *E. coli* SM10 | Cloning and mating strain, *thi-1, thr leu tonA lacY supE recA::RP4-2-Tc::Mu pir (Km<sup>R</sup>)* | 33 |
| **Plasmid** |  |  |
| pEX18Tc | multi-host suicide vector, Tet<sup>R</sup>, *oriT*, *sacB*, *lacZα*, MCS from pUC18 | 34 |
| pJLM2-WTA | pEX18Tc with *aguA* from PA14 WT | This study |
| pJLM2-*ΔaguA* | pEX18Tc with mutant *aguA* from MNPA05 | This study |
| pJLM2-*ΔspeA* | pEX18Tc with *speA* knockout construct | This study |

### Agmatine Biosensor assay

Clinical isolates were screened for agmatine accumulation using a modified bioassay described previously (31). Briefly, an agmatine reporter strain defective in agmatine metabolism (PA14 *aguA:gm Δagu2ABCA’ ΔspeA aguRB*-lux) was grown overnight in LB to stationary phase and diluted in fresh LB to a final OD_600_ of ~0.2. Spent supernatants from the reporter strain were used to generate a standard curve, whereby exogenous agmatine (Sigma) was added at a concentration of 200 µM, followed by serial dilution to 0.2 µM. 100 µL of each standard was added, in triplicate, to a 96-well microtiter plate. Stationary phase cultures (n=3) of *P. aeruginosa* clinical isolates (92 strains in total) were then centrifuged at 10,000 x g for 5 min, and 100µL of spent supernatant was added in triplicate to plates containing the standards. Finally, 100 µL of the reporter strain was added to each well, plates were covered with a Titer Top seal (Diversified Biotech) to prevent evaporation and incubated at 37°C for 3h. Luminescence was determined using a BioTek Synergy H1 plate reader and normalized to culture density (OD_600_) of the reporter strain.

### PCR amplification and sequencing of *aguA*

AccuPrime GC-Rich DNA polymerase kit (Invitrogen) and primers aguAF and aguAR (Table S1) were used to amplify *aguA* from each clinical isolate. PCR products were gel purified using QIAquick Gel Extraction kit (Qiagen) and sent to Functional Biosciences (Madison, WI) or Eurofins Genomics (Louisville, KY) for sequencing.

### Plasmid construction and genetic manipulation

*aguA* from *P. aeruginosa* PA14 (WT gene) was PCR amplified and cloned into HindIII/SmaI-digested pEX18Tc using primers aguAFHindIII and aguAR forming pJLM2-WTA. To restore agmatine catabolism in MNPA04, MNPA05, MNPA06, CHB47-6 and CHB54-11, pJLM2-WTA was transformed into *E. coli* SM10 and mobilized into the clinical isolates. Recombinants (containing the native *aguA* and intact WT *aguA* gene from strain PA14) were selected on LB agar containing irgisan (25 µg/mL) and tetracycline (25-200 µg/mL, depending on the isolate). Colonies were then transferred to LB agar containing 200 µg/mL tetracycline, grown overnight, and patched to 5% sucrose plates. Genomic DNA from colonies that grew on sucrose was then screened by PCR and sequenced using primers aguAF and aguAR1 to confirm recombination. *speA* deletion mutants were generated by cloning and ligating a *speA* knockout construct (31) into HindIII/EcoRI-digested pEX18Tc using primer pair speAF and speAR, generating pJLM2-ΔspeA. This plasmid was then transformed into *E. coli* SM10 and mobilized into *P. aeruginosa* as described above.

### Antibiotic susceptibility assay

Minimum inhibitory concentrations of ceftazidime hydrate (Sigma), doxycycline hyclate (Sigma), gentamicin sulfate (Amresco), tobramycin (Amresco), colistin sulfate (Santa Cruz Biotechnology), and piperacillin sodium (Gold Biotechnology) were determined using a microtiter plate-based assay. Tazobactam (10 µg/mL) was also added to the piperacillin treatments. Briefly, bacterial cultures were grown overnight in LB, diluted to an OD_600_ of ~0.2, and further diluted 100-fold in MHB-2. 96-well plates were then prepared with 100 μL of MHB-2 containing two-fold dilutions of antibiotics, followed by the addition of 100 μL of diluted overnight cultures to each well. Cell-free wells for each antibiotic and MHB-2 alone (no antibiotics) were used as controls. Plates were sealed with Titer Top adhesive and placed at 37°C for 24h. Cell density was measured spectrophotometrically (OD_600_) using in a BioTek Synergy H1 plate reader. Strains were tested in triplicate using four technical replicates per plate. Bacterial densities versus six, 2-fold dilutions of each antibiotic were then fit with a non-linear regression using a four-parameter logistic function to determine IC50 values and were compared using one-way Kruskal-Wallace ANOVA. All statistical analyses were performed in Graphpad Prism v.6.0.

### Acute pneumonia mouse model

Overnight cultures of MNPA04 and MNPA04+*aguA* were grown in LB, washed once in PBS, and resuspended to an OD_600_ of ~1.0. Two groups (n=10) of female BALB/c mice (Jackson Laboratories), aged 8 weeks, were anesthetized using isoflurane (3% at 3L/min) and challenged intratracheally with 100µL (1 x 10^8^ c.f.u.) of culture using a 1mL Hamilton syringe and a 22G × 1.25” catheter (Terumo Medical). After 24h, mice were sacrificed via CO_2_ asphyxiation and cervical dislocation, followed by bronchoalveolar lavage using 2mL of sterile PBS. Lavage fluid (BAL) was then serially diluted ten-fold and plated on LB agar + irgisan to quantify *P. aeruginosa* load. A 100 µL aliquot of BAL was centrifuged in a Cytospin column (ThermoFisher), stained for differential white cell counts, and neutrophils were quantified using a hemocytometer. Neutrophil counts and BAL colony forming units were compared using unpaired two-tailed t-tests with Welch’s correction. Animal work followed guidelines of the Association for Assessment and Accreditation of Laboratory Animal Care (AAALAC) and was approved by the UMN Institutional Animal Care and Use Committee (#1212-30122A).

### Epithelial cell culture model

CF-DHBE bronchial epithelial cells were seeded into 48-well cell culture plates (Costar) coated with bovine collagen at 5×10^4^ cells per well in 250µL of BEGM. Cells were incubated at 37°C in humidified air with 5% CO_2_. Media was replaced every 48h until cells were confluent on day 10 post-seeding. Prior to treatment, cell cultures were rinsed twice with 250µL starvation medium (MEM + 0.5% FBS). Cells were then stimulated by adding 10-fold dilutions of agmatine (0.1 – 100 µM)(Sigma), 100 ng/mL of LPS from *E. coli* (Sigma), or agmatine plus LPS in starvation medium to each well. Media-only and media+LPS were used as controls. Treated cells were incubated for an additional 24h, followed by collection of supernatants that were frozen at -80°C for downstream analysis. Supernatants were assayed using the IL-8 Human ProcartaPlex^®^ Simplex Kit (Thermo) on the Luminex Magpix instrument according to manufacturer’s protocol. Standards were run in duplicate and test samples were run in triplicate on the same plate. Cytokine concentrations were compared using a non-parametric Friedman’s test with Dunn’s multiple comparison post-test.

## RESULTS

### Agmatine accumulation among *P. aeruginosa* CF isolates

Given the elevated concentrations of agmatine detected within expectorated sputum and the identification of overproducing strains (26), we first tested our hypothesis that agmatine accumulation (herein referred to as “hyperproduction”) is a common phenotype among *P. aeruginosa* CF isolates. To evaluate its frequency, we used a previously described biosensor assay (31) to quantify agmatine production among a strain library comprising 92 CF airway isolates. Isolates were grown to stationary-phase, and supernatants were added to an agmatine-sensitive bioluminescent reporter derivative of *P. aeruginosa* PA14 (31). Luminescence was then quantified after 3h and normalized to culture density of the reporter strain (Fig. 2). Of ninety-two isolates tested, five (5.4%) were found to produce agmatine above the detectable threshold of ~1 µM, including three that were identified previously (26). These concentrations were equal to or above those produced by PA14Δ*aguA*Δ*agu2Aagu2A’*, an agmatine hyperproducing lab strain used in the development and validation of the assay (31).

**Figure 2.**
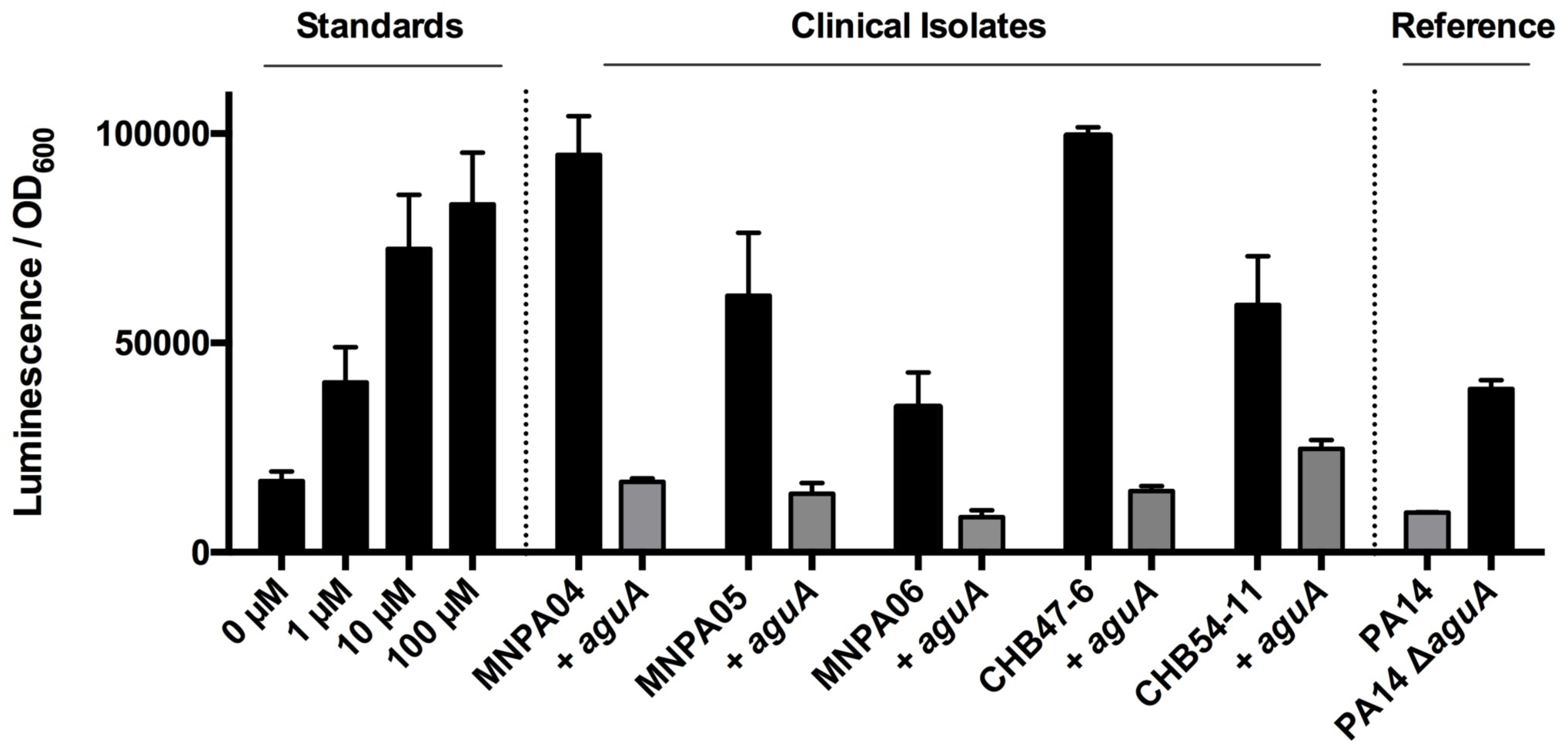
Agmatine biosensor assay of *P. aeruginosa* clinical isolates. Agmatine hyperproduction was detected in 5/92 clinical isolates (black bars). Data shown are the mean (n=3) luminescence normalized to biosensor culture density. Chromosomal complementation with intact *aguA* from PA14 restores agmatine metabolism (grey bars).

Since *aguA* encodes a deiminase that catalyzes the conversion of agmatine to N-carbamoyl putrescine (Fig. 1), we predicted that mutations at this locus were responsible for agmatine accumulation. To test this, *aguA* was PCR amplified from each isolate and sequenced. Interestingly, MNPA04, MNPA05, and MNPA06 each harbored an identical 11 base pair deletion in *aguA* (Fig. S1), which may be indicative of patient-to-patient transmission of an agmatine-hyperproducing common ancestor. CHB47-6 and CHB54-11 also harbored 8-bp and 14-bp deletions, respectively, at the 3’ end of *aguA* (Fig. S1). Each of these deletions results in a frameshift mutation in the downstream catalytic site of the deiminase. Notably, none of the five hyperproducing isolates contained the alternate *agu2ABCA’* operon (encoding two additional agmatine deiminases) found in ~20% of *P. aeruginosa* isolates (32), suggesting that mutations identified in *aguA* were responsible for the agmatine hyperproduction phenotype. Indeed, when an intact copy of *aguA* was cloned from strain PA14 and mobilized via allelic exchange into the *aguA* mutant isolates, agmatine catabolism was restored as determined by the biosensor assay (Fig. 2).

### Agmatine hyperproduction confers increased tolerance to cationic antibiotics

The isolation of *aguA* mutants from multiple subjects raised the question of whether agmatine hyperproduction is another example of a loss-of-function phenotype that confers a fitness advantage. For example, does agmatine decrease susceptibility to positively charged antibiotics that destabilize the Gram-negative outer membrane? This question was raised, in part, based on prior reports of exogenously added spermidine and putrescine conferring protection to *P. aeruginosa* from polymixin B (29, 35). This was likely due to increased stability of lipopolysaccharide (LPS) via the polycationic charge of the polyamines (29). Though agmatine is a pre-polyamine, it also harbors a dipositive charge, and we hypothesized that its production by the *aguA-* clinical isolates confers a similar protective effect.

Using a microtiter plate-based assay, the minimum inhibitory concentration and IC50 (antibiotic concentration needed to halve the bacterial density at 24h) (36) of three commonly used CF antibiotics with a polycationic charge (colistin, tobramycin, and gentamicin) were determined for MNPA04 and its isogenic *aguA*+ strain. To ensure that any observed phenotype was specific for agmatine accumulation (and not due to a defect in arginine catabolism), we also generated a MNPA04Δ*speA* mutant lacking the arginine decarboxylase (SpeA) required for the first dedicated step of the pathway (see Fig. 1). While there was no difference in MIC for any of the compounds tested (data not shown), there was a significant increase in IC50 for each antibiotic versus the agmatine hyperproducer relative to the complemented *aguA*+ strain (Fig. 3A, Table 2). This phenotype was not observed in MNPA04Δ*speA*, suggesting that the observed decreases in antibiotic susceptibility were a direct result of agmatine accumulation.

**Table 2.**
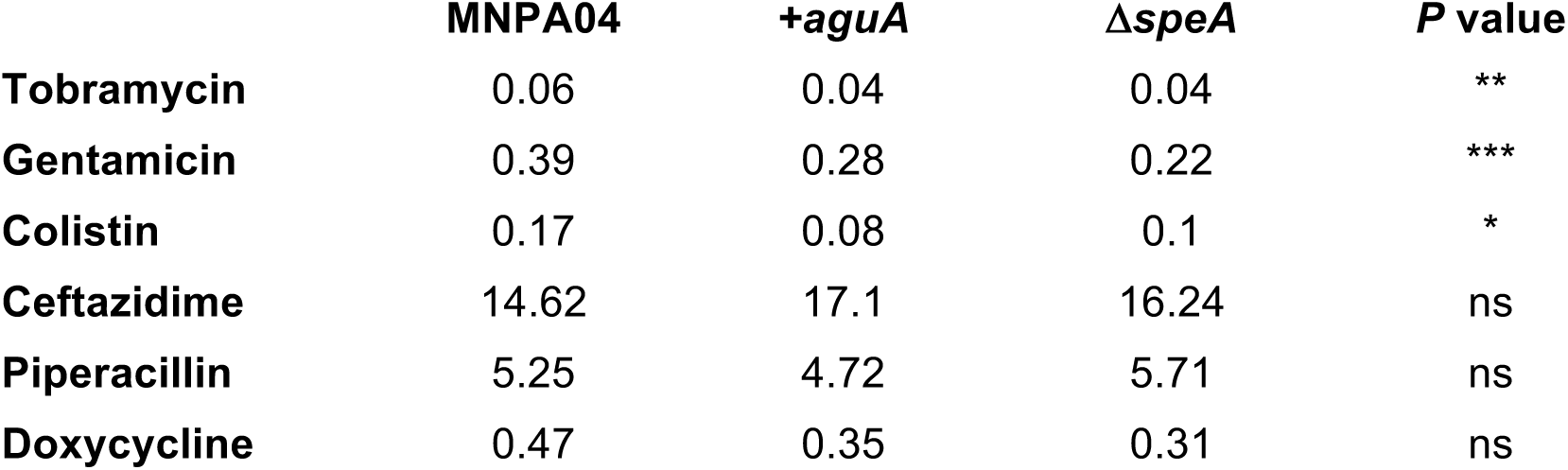
IC50 values (µg/mL) for the agmatine hyperproducing clinical isolate (MNPA04) of *P. aeruginosa*.

**Figure 3.**
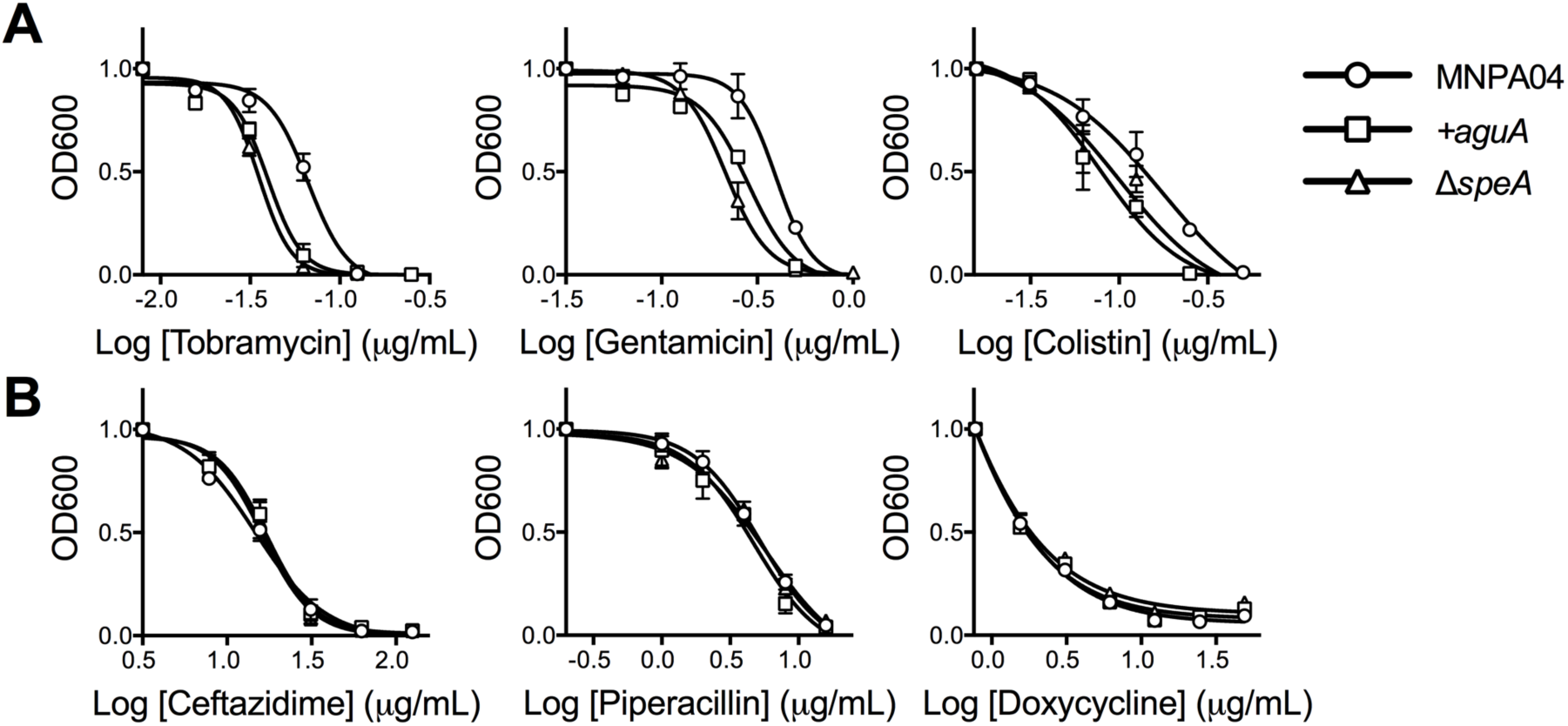
Agmatine hyperproduction confers increased tolerance to cationic antibiotics. (A) Agmatine hyperproduction by clinical isolate MNPA04 confers increased tolerance to the positively charged tobramycin, gentamicin and colistin. Susceptibility is restored when complemented with *aguA* or by deletion of *speA*. (B) No differences in susceptibility were observed for antibiotics with a neutral or negative charge. IC50 values (the concentration of drug required to halve the bacterial density) were determined by non-linear regression analysis using a four-parameter logistic function (see Fig. S2).

As demonstrated previously for spermidine and putrescine (29,35), we predicted that the dipositive charge of agmatine impeded the interaction of cationic antibiotics with the negatively charged bacterial cell surface. To test this, we treated MNPA04, MNPA04+*aguA* and MNPA04Δ*speA* with three additional CF antibiotics that carry a net neutral or negative charge at physiological pH; ceftazidime, piperacillin, and doxycycline (Fig. 3B, Table 2). As predicted, there were no significant changes in IC50 for the neutral or electronegative compounds, supporting the hypothesis that agmatine hyperproduction decreases *P. aeruginosa* susceptibility to cationic antibiotics due its dipositive charge.

Interestingly, resistance phenotypes were strain-dependent; when tobramycin, gentamicin and colistin were tested against other mutant isolates identified in our screen, agmatine conferred increased resistance in just three of five strains (Table S2). Strain-to-strain variability was not observed for the neutral and negatively charged compounds. Notably, the two isolates for which increased resistance to the aminoglycosides (tobramycin and gentamicin) was not found also had observable phenotypes that may also contribute to antibiotic tolerance – pyomelanin hyperproduction (MNPA06) and mucoidy (CHBPA54-11)(Fig.S3)(37–39). It is possible that these and/or other phenotypic characteristics affect the agmatine-antibiotic interaction that was observed for MNPA04, MNPA05, and CHB47-6. Nevertheless, the data suggest that deletions in *aguA* and the consequent accumulation of agmatine may be beneficial for some *P. aeruginosa* clinical isolates.

### Mouse pneumonia model

Murine models have revealed both a beneficial and detrimental role of agmatine in reducing acute lung injury through modulation of inflammatory cytokines (26, 40). However, agmatine has also been shown to reduce macrophage TNF-α and MIP-2 response to bacterial LPS (26). This led to our hypothesis that in a murine acute pneumonia model, the airway inflammatory response to agmatine hyperproducing clinical isolates (*aguA-*) of *P. aeruginosa* would also be dampened relative to *aguA*+ strains. To test this hypothesis, BALB/c mice were challenged intratracheally with either MNPA04 or MNPA04+*aguA* for 24h, followed by quantification of neutrophil recruitment into the airways. On average, MNPA04 recruited 40% fewer neutrophils than the *aguA*+ strain (*P =* 0.002, Figure 4A). To determine whether this decrease was a result of impaired bacterial growth *in vivo*, *P. aeruginosa* colony forming units in BAL were also quantified. Consistent with previous studies (41), a three-log reduction in colony forming units was observed relative to the inoculum after 24h, yet there was no significant difference between treatment groups (*P*=0.48, Figure 4B). These data suggest that despite a reduction of initial bacterial load, neutrophil recruitment is impaired by *P. aeruginosa* agmatine hyperproduction.

**Figure 4.**
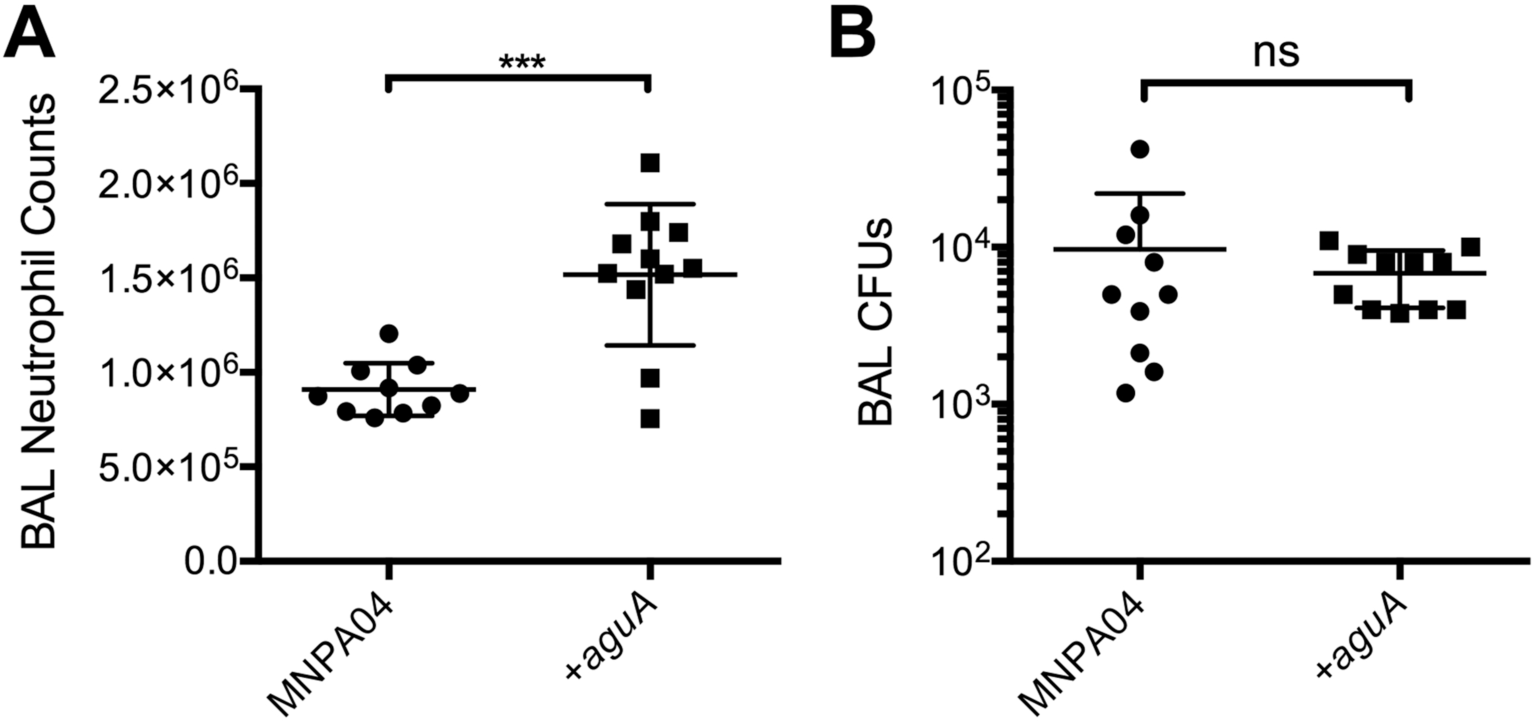
Agmatine hyperproduction by *P. aeruginosa* affects neutrophil recruitment to the airways. Mice were subjected to a 24h airway infection with either MNPA04 (*aguA-*) or a MNPA04+*aguA* strain. (A) Neutrophils counts from the BAL fluids were significantly reduced (P=0.0002) when the mice were challenged with an agmatine overproducing *P. aeruginosa* isolate. (B) Bacterial loads were comparable between treatments, ruling out *in vivo* growth differences. Each data point represents an individual treatment. Means were compared using an unpaired t-test using Welch’s correction for variance.

### Epithelial cell model

Neutrophil recruitment to a site of infection is predominately mediated by the pro-inflammatory cytokine IL-8 in response to pathogen associated molecular patterns (PAMPs)(42–44). Therefore, we speculated that the observed decrease in neutrophil recruitment *in vivo* was a result of *P. aeruginosa-*derived agmatine diminishing MIP-2 (the murine version of IL-8) production. To test this, primary CF-DHBE airway epithelial cells (AECs) that were homozygous for ΔF508 in the gene encoding the cystic fibrosis transmembrane regulator (CFTR) protein were treated with lipopolysaccharide (100 ng/mL) in the presence of increasing concentrations of agmatine (1-100 µM), followed by quantification of IL-8 production in the AEC culture supernatant (Fig. 5). Relative to AECs treated with LPS treatment alone (2.8 ng/mL), IL-8 levels were significantly reduced for AECs treated with LPS+100µM agmatine (1.2 ng/mL; *P =* 0.02). These IL-8 levels were comparable to media and media+agmatine controls, suggesting that agmatine can reduce the pro-inflammatory response of AECs to bacterial infection. These data corroborate our *in vivo* observations of the host response to agmatine hyperproducing clinical isolates of *P. aeruginosa*. Furthermore, they support our hypothesis that agmatine may confer an *in vivo* persistence advantage for *P. aeruginosa* in the context of CF lung infection.

**Figure 5.**
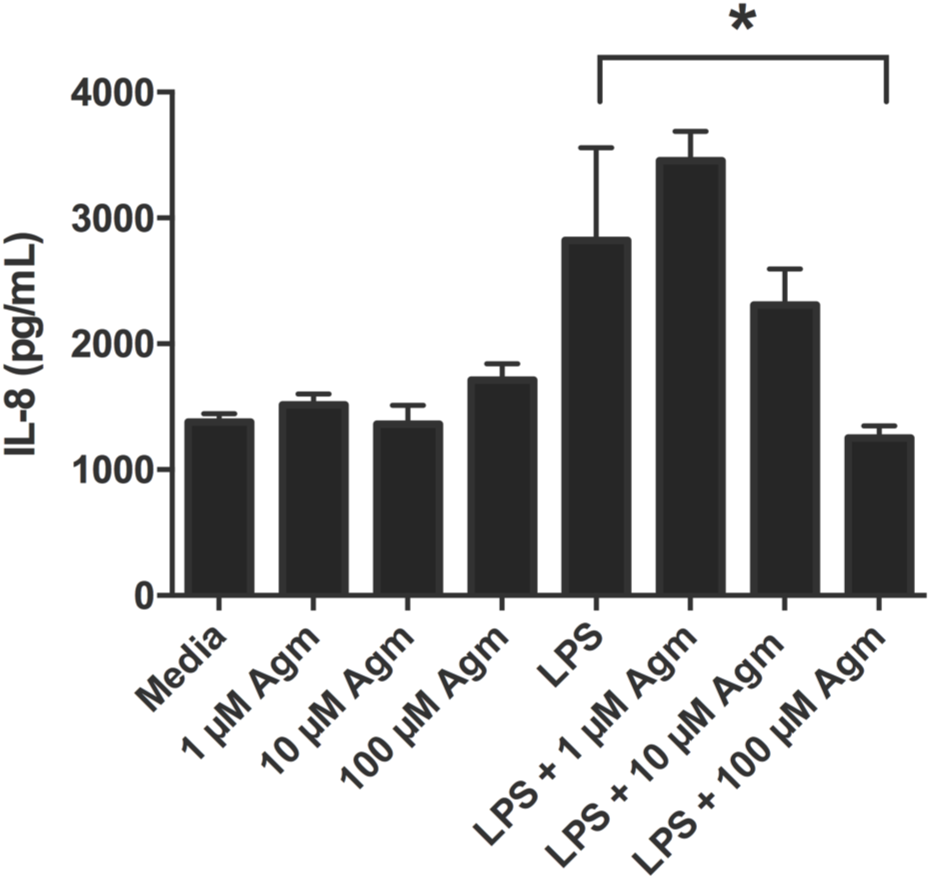
Airway epithelial cell response to agmatine and LPS. CF-DHBE primary epithelial cells were treated with either agmatine or LPS alone, or agmatine with LPS for 24 hours. Agmatine diminishes the IL-8 response to LPS challenge. Data shown are means + SD (n=3) and were compared using a one-way Friedman’s test (*P=*0.0061) with Dunn’s multiple comparison post-test. IL-8 levels were significantly reduced for AECs treated with LPS+100µM agmatine relative to LPS alone (1.2 ng/mL; *P =* 0.02).

## DISCUSSION

Polyamines are low molecular weight polycations ubiquitously found in all living cells. Owing to their regularly-spaced positive charges, these metabolites play critical roles in cell growth and proliferation, transcription and translation, signal transduction, ion transport and other cellular processes (45, 46). An important precursor to polyamine formation, agmatine, is a polycationic intermediate of the arginine decarboxylase pathway for which diverse roles have also evolved. In mammals, for example, agmatine exerts its effect at multiple targets, including matrix metalloprotease and NADPH oxidase regulation, neuro- and immunomodulation, and nitric oxide synthesis (26,47–49). Recently, agmatine was also proposed as an environmental trigger for *P. aeruginosa* biofilm formation (32), as growth of strain PA14 in the presence of exogenous agmatine showed a dose-dependent increase in biofilm development, while deletion of both *aguA* and *agu2ABCA’* (encoding redundant agmatine deiminases that convert agmatine into N-carbamoylputrescine, Fig. 1) showed a significant increase in biomass relative to WT. Here we demonstrate that *aguA-* mutants of *P. aeruginosa* that hyperproduce agmatine are frequently isolated from the airways of CF patients. Moreover, we present evidence that agmatine accumulation in the extracellular milieu may confer multiple selective advantages for persistence of *P. aeruginosa in vivo*.

Smith et al. (15) demonstrated that during chronic airway infection, *P. aeruginosa* undergoes adaptive evolution through genomic variation, and that a large number of genes are targets for mutation. Mutations in the gene encoding the key quorum-sensing regulator, *lasR*, are especially common, and lead to defects in the production of key virulence effector compounds (12). Loss-of-function adaptations in *gacS*, *retS, mutS, mucA* and *ampR* (50–53) also increase in prevalence as infections progress and reflect the transition of *P. aeruginosa* from an acute to chronic infection phenotype. Many of these examples are mutated in only a small fraction of infections (15), which is consistent with the prevalence of *aguA* mutations (~5%) among our isolate library. While this recovery rate is not nearly as high as QS-deficient (*lasR*), mucoid (*mucA)* or hypermutator (*mutS)* phenotypes (12, 54, 55), parallel evolution of agmatine hyperproduction in *P. aeruginosa* derived from multiple patients suggests positive selection at the *aguA* locus. Since our isolate library was derived from de-identified sputum samples for which clinical data were not available, it is unclear whether agmatine hyperproduction is associated with the transition to chronicity. It is also not known whether *aguA* mutations are selected for by the *in vivo* microenvironment, or if *aguA-* environmental isolates are better poised for airway colonization. Future studies aimed at a larger patient cohort with detailed clinical data will help address these questions.

The observed decrease in susceptibility of *aguA-* strains to both aminoglycosides (gentamicin, tobramycin) and polymixins (colistin) suggests that the observed effect is due to a shared characteristic of the antibiotics (*e.g*. their positive charge). While primary cellular targets differ, the cationic nature of both drug classes is thought to increase outer membrane (OM) permeability to lysozyme and hydrophobic compounds (56,57). The initial action of aminoglycosides has been shown to cause disruption of salt-bridges between adjacent LPS molecules, disrupting the normal OM packing order and allowing the compound to then enter the cell and inhibit protein translation (57,58). Similarly, polyamines carry a polycationic charge that facilitates binding to LPS. However, owing to their small size, polyamines do not disrupt outer membrane packing or increase OM permeability on their own (58). In fact, the polyamines spermidine and putrescine have been shown to protect Gram-negatives against antimicrobials and oxidative stress, in addition to stabilizing spheroplasts and protecting them against lysis (29, 35, 59, 60). Given that Δ*speA* mutants (defective in arginine decarboxylation) showed no resistance phenotype and that neutrally charged antibiotics were not affected by *aguA* mutations, we propose that agmatine also confers resistance to *P. aeruginosa* due to its dipositive charge.

The observed strain-to-strain variability suggesting that agmatine-antibiotic inhibition can be influenced by additional factors was not unexpected. Alginate, for example, has been shown to restrict the diffusion of aminoglycosides (cationic), but not β-lactams (neutral), through mucoid biofilms of *P. aeruginosa* clinical isolates (61). Elevated concentrations of salts increased aminoglycoside diffusion, suggesting that electrostatic interactions between cationic compounds and extracellular polymers can also impact their efficacy (61,62). It remains to be determined whether variations in LPS antigens or the production of the extracellular polysaccharides pel and psl that differ in charge and vary among clinical isolates, also impact susceptibility to antibiotic challenge in the presence of agmatine (63,64). Likewise, pyomelanin, an electronegative pigment that is overproduced due to mutations in *hmgA*, exhibits humic-type properties that allows for chelation of soluble cations (65). It is possible that (in the case of MNPA06) its polyanionic charge may impact cell surface interactions with polyamines and antibiotics. We are currently using whole genome sequencing and phenotypic assays of the *aguA-* mutants to understand how these and other phenotypes (including biofilm formation) influence the protective effects of agmatine for *P. aeruginosa*.

CF airway disease is characterized by chronic, neutrophil-dominated inflammation in response to bacterial infection. However, *P. aeruginosa* can also use numerous strategies to evade detection and eradication by the immune system. These include formation of biofilms that prevent phagocytic clearance, altered expression of PAMPs such that detection by host immune receptors and downstream signaling is minimized, and direct interference with host signaling through effector molecules (66–67). In the case of agmatine, impaired recruitment of neutrophils is an example of the latter. More specifically, we favor the interpretation that the dipositive charge of agmatine prevents binding of LPS by LPS-binding protein (LBP) and the downstream production of pro-inflammatory cytokines (*e.g*. IL-8). These data are consistent with previous reports of diminished MIP-2 production in murine peritoneal macrophages in response to LPS in the presence of agmatine (26). Given that cationic antimicrobial peptides have been shown to block LPS-LBP interactions and macrophage cytokine production (68), our data support a similar mechanism for agmatine.

It has been shown that agmatine is capable of both immune activation and inhibition depending on the presence and dose of co-stimulatory molecules (26). This dichotomy may explain, at least in part, contrasting phenotypes between the lung adapted isolate MN004 reported here, and those found for the virulent burn wound isolate PA14 (26) which is known to produce an array of immunostimulatory exoproducts (*e.g*. pyocyanin). Similarily, the acute pneumonia model used here revealed no difference in bacterial load between *aguA-* and *aguA+* variants of MN004, whereas an agmatine hyperproducing mutant of PA14 showed a significant *in vivo* growth defect relative to the wildtype (26). These strain-to-strain variations, similar to those observed for antibiotic tolerance, underscore the potential combinatorial effect of multiple phenotypes on agmatine-mediated interactions of *P. aeruginosa* with its growth environment.

Despite these variations, agmatine hyperproduction has potential consequences for long-term *P. aeruginosa* persistence *in vivo*. For example, increased tolerance to positively charged therapeutics would confer a significant growth advantage, particularly in mucus-plugged, diffusion-restricted airways with sub-inhibitory antibiotic concentrations. Further, by accumulating *aguA* mutations, agmatine hyperproduction would lead to an impaired ability for the host to mount an adequate reaction aimed at pathogen clearance, in turn creating an environment that facilities adaptation and chronic persistence. Further experiments using a chronic bead model of infection will be used to directly test this possibility. Either or both of these advantages, coupled with agmatine-mediated biofilm formation reported previously (32), represent a multifactorial basis for the positive selection for *aguA* mutants and may be a contributing factor to their success in the CF lung environment.

## ACKNOWLEDGEMENTS

This work was supported by National Institutes of Health K08 HL105466 (BJW) and a Pathway to Independence Award (R00HL11462) awarded to RCH. JLM was supported by a National Institutes of Health Lung Sciences T32 fellowship (#2T32HL007741-21) awarded through the National Heart Lung & Blood Institute. We thank Gary Dunny, Yinduo Ji and Scott O’Grady (University of Minnesota) for their guidance, and members of the Hunter lab for their expertise and critical review of the manuscript.

